# Allele mining unlocks the identification of RYMV resistance genes and alleles in African cultivated rice

**DOI:** 10.1101/2020.01.22.914143

**Authors:** Hélène Pidon, Sophie Chéron, Alain Ghesquière, Laurence Albar

**Affiliations:** DIADE, Univ. Montpellier, IRD, Montpellier, France

**Keywords:** rice, *Oryza glaberrima*, RYMV, resistance gene, CPR5

## Abstract

**Background:** *Rice yellow mosaic virus* (RYMV) is a major rice pathogen in Africa. Three resistance genes, i.e. *RYMV1*, *RYMV2* and *RYMV3,* have been previously described. *RYMV1* encodes the translation initiation factor eIF(iso)4G-1 and the best candidate genes for *RYMV2* and *RYMV3* encode a homolog of an *Arabidopsis* nucleoporin (CPR5) and a nucleotide-binding domain and leucine-rich repeat containing domain (NLR) protein, respectively. High resistance is very uncommon in Asian cultivated rice (*Oryza sativa*), with only two highly resistant accessions identified so far, but it is more frequent in African cultivated rice (*Oryza glaberrima*).

**Results:** Here we report the findings of a resistance survey in a reference collection of 268 *O. glaberrima* accessions. A total of 40 resistant accessions were found, thus confirming the high frequency of resistance to RYMV in this species. We analysed the variability of resistance genes or candidate genes in this collection based on high-depth Illumina data or Sanger sequencing. Alleles previously shown to be associated with resistance were observed in 31 resistant accessions but not in any susceptible ones. Five original alleles with a frameshift or untimely stop codon in the candidate gene for *RYMV2* were also identified in resistant accessions. A genetic analysis revealed that these alleles, as well as T-DNA insertions in the candidate gene, were responsible of RYMV resistance. All 40 resistant accessions were ultimately linked to a validated or candidate resistance allele at one of the three resistance genes to RYMV.

**Conclusion:** This study demonstrated that the *RYMV2* resistance gene is homologous to the *Arabidopsis CPR5* gene and revealed five new resistance alleles at this locus. It also confirmed the close association between resistance and an amino-acid substitution in the leucine-rich repeat of the NLR candidate for *RYMV3*. We also provide an extensive overview of the genetic diversity of resistance to RYMV in the *O. glaberrima* species, while underlining the contrasted pattern of diversity between *O. glaberrima* and *O. sativa* for this trait. The different resistance genes and alleles will be instrumental in breeding varieties with sustainable field resistance to RYMV.

## Background

*Oryza sativa*, domesticated in Asia, is cropped in almost all rice-growing areas worldwide. However, an independent rice domestication process occurred in Africa, which gave rise to the cultivated species *Oryza glaberrima* [1, 2]. The more productive *O. sativa* species was introduced in East Africa more than 1,000 years ago and in West Africa in the 16th century, and has progressively supplanted *O. glaberrima*. Breeding initiatives over the last 60 years have essentially concerned *O. sativa* varieties and have further widened the gap in yield potential between varieties of the two species. Nonetheless, *O. glaberrima* has specific traits of interest and adaptation to local stresses, such as drought, iron toxicity, infertile soils and weed competition [3, 4]. This rich source of gene diversity is of substantial breeding interest to increase rice yield in a setting of global warming and reduced inputs. *O. glaberrima* was thus introduced in breeding programs [5, 6] leading for instance to the New Rice for Africa (NERICA) varieties, that resulted from *O. sativa* x *O. glaberrima* interspecific crosses and were successfully disseminated in the 2000s [7, 8].

The *Rice yellow mottle virus* (RYMV) is endemic to Africa and responsible for significant rice crop losses in irrigated or lowland areas [9]. High resistance appears to be very uncommon in *O. sativa*, with only two highly resistant varieties identified so far [10, 11], whereas 31 highly resistant *O. glaberrima* accessions have been reported [12, 13]. Moreover, while the two *O. sativa* resistant varieties share the same allele of the *RYMV1* resistance gene, which encodes a translation initiation factor, at least three different *RYMV1* resistance alleles evolved independently in *O. glaberrima* [12, 14]. These results suggest that *O. glaberrima* diversity for this trait would be particularly useful for rice breeding.

Two additional resistance genes, i.e. *RYMV2* and *RYMV3,* have been mapped in *O. glaberrima* species. *RYMV2*-mediated resistance is associated with a 1 bp deletion, leading to a null allele of an homolog of the *Arabidopsis constitutive expression of pathogenesis related protein-5 (CPR5)* nucleoporin gene in both a bi-parental mapping population and a collection of *O. glaberrima* accessions [13]. In *Arabidopsis,* the *CPR5* nucleoporin gene is involved in the regulation of defense mechanisms and senescence [15, 16]. Considering the sequence homology and in line with previous studies [13, 17], the candidate gene for *RYMV2* is hereafter referred to as *CPR5-1,* although its nucleoporin role and involvement in defense mechanisms has yet to be documented in rice. More recently, a gene of the nucleotide binding domain and leucine-rich repeat gene (NLR) superfamily was pinpointed as the best candidate for the *RYMV3* dominant resistance gene [18]. This gene is hereafter referred to as *NLR_RYMV3_*. Resistance is associated with a single amino-acid substitution in the leucine-rich repeat (LRR) domain of the protein, which is known to be involved in the pathogen recognition specificity.

Here we describe the diversity of RYMV resistance genes or candidates in one of the most documented *O. glaberrima* collections, which covers the geographical distribution of the species and includes 165 fully sequenced accessions [19, 20]. We also validated the candidate gene for *RYMV2* using natural variants identified in *O. glaberrima* diversity and *O. sativa* T-DNA mutants.

## Results

### Screening for resistance to RYMV in a collection of *O. glaberrima* accessions

Thiemele et al. [12] and Orjuela et al. [13] screened 120 accessions of the *O. glaberrima* collection described in Orjuela et al. [19] for resistance to RYMV and found 31 highly resistant accessions. In the present study, these 31 accessions and 148 additional ones from the same collection were phenotyped for resistance by double antibody sandwich enzyme-linked immunosorbent assay (DAS-ELISA) on a set of four plants per accession. The same virus isolate as that reported in Thiemele et al. [12] and Orjuela et al. [13] was used. Of the 31 accessions previously reported as being resistant, we confirmed the resistance of 28, while three were susceptible, presumably because of between seed stocks heterogeneity. All four plants of most of the 148 newly tested accessions were clearly susceptible. However, highly resistant plants were observed in 12 accessions for which the high resistance phenotype was confirmed in additional plants when seeds were available (Table 1). For eight of those, up to a third of the plants multiplied the virus, suggesting incomplete resistance or possible resistance-breaking events, as previously reported [17,18,21]. However, the rate of susceptible *vs*. resistant plants was not significantly different than observed on the Tog7291 accession carrying the *RYMV2* major gene (Fisher exact test, p>0.05) and these accessions were thus considered resistant. Finally, the accessions identified as being resistant in this study were: Og26, Og111, Og133, Og183, Og213, Og256, Og406, Og423, Og447, Og452, Og491 and Og498 (Table 1; Additional file 1: Table S1). A total of 40 accessions out of 268 were therefore highly resistant to the BF1 isolate of RYMV.

**Table 1.** Phenotyping of O. glaberrima accessions for RYMV resistance. Only accessions identified as resistant in this study are listed in this table. Resistance was evaluated based on ELISA tests performed on individual plants after mechanical inoculation with the BF1 isolate of RYMV. The first screening was performed on a set of four plants and confirmed, when seeds were available, in additional screening experiments. Only accessions for which high resistance was observed are listed. Tog5681 and Tog7291, carrying resistance alleles on *RYMV1* and *RYMV2* genes, respectively, were used as resistance controls and Og82, Og431 and CG14 were used as susceptible controls. The total number of resistant and susceptible plants and the percentage of resistant plants are indicated.

| Accessions | Resistant | Susceptible | Resistance rate (%) |
| --- | --- | --- | --- |
| Susceptible controls | 2 | 22 | 8 |
| Tog5681 | 8 | 0 | 100 |
| Tog7291 | 10 | 2 | 83 |
| Og26 | 18 | 0 | 100 |
| Og111 | 12 | 0 | 100 |
| Og133 | 11 | 6 | 65 |
| Og183 | 16 | 2 | 89 |
| Og213 | 9 | 4 | 69 |
| Og256 | 14 | 4 | 78 |
| Og406 | 15 | 1 | 94 |
| Og423 | 4 | 0 | 100 |
| Og447 | 14 | 2 | 87 |
| Og452 | 16 | 0 | 100 |
| Og491 | 17 | 5 | 77 |
| Og498 | 16 | 2 | 89 |

### Allele mining in RYMV resistance genes or candidates

Among resistant *O. glaberrima* accessions, previous results indicated that 12 have a resistance allele on the *RYMV1* gene [12], 7 have an allele associated with *RYMV2*-mediated resistance on the *CPR5-1* gene [13], 1 has a resistance allele on *RYMV3*, for which *NLR_RYMV3_* is a candidate [18], while the Tog5672 accession carries a resistance allele on both *RYMV1* and *RYMV3* [18]. The genes or alleles responsible for resistance in the 19 remaining accessions were unknown.

Polymorphisms in the *RYMV1* gene, and in the *RYMV2* and *RYMV3* candidate genes, the *CPR5-1* gene and *NLR_RYMV3_*, respectively, were analyzed in 165 accessions for which the full genome sequence was available [1,20,22]. For the nine resistant accessions for which the full genome sequence was not available, the partial or complete sequence of the target genes were obtained from Thiemele et al. [12] or by Sanger sequencing of polymerase chain reaction (PCR) fragments.

### Allele mining in *RYMV1*

A total of ten single nucleotide polymorphisms (SNPs) or small insertions/deletions (indels), defining nine different haplotypes, were detected in the *RYMV1* gene (Additional file 1: Table S2). The three most frequent haplotypes at the nucleotidic level represented 83% of the accessions and corresponded to the protein variant of the CG14 accession, while the others were detected in less than 5% of the accessions. Five mutations – three SNPs and two indels – were located in the exons and all resulted in amino-acid changes (Figure 1A). One of them that caused a single amino-acid substitution (P541L) was present in susceptible accessions. The others, which were previously described as characterizing *rymv1-3* (R322_D324del, S576N), *rymv1-4* (E321K) and *rymv1-5* (K312_G315delinsN) resistance alleles [12, 14], were associated with resistance in the full collection. Fifteen resistant accessions carried those resistance alleles (Table 2; Additional file 1: Table S1), including two accessions identified as resistant in this study, Og208 and Og423, which carried alleles *rymv1-3* and *rymv1-4*, respectively.

**Figure 1.**
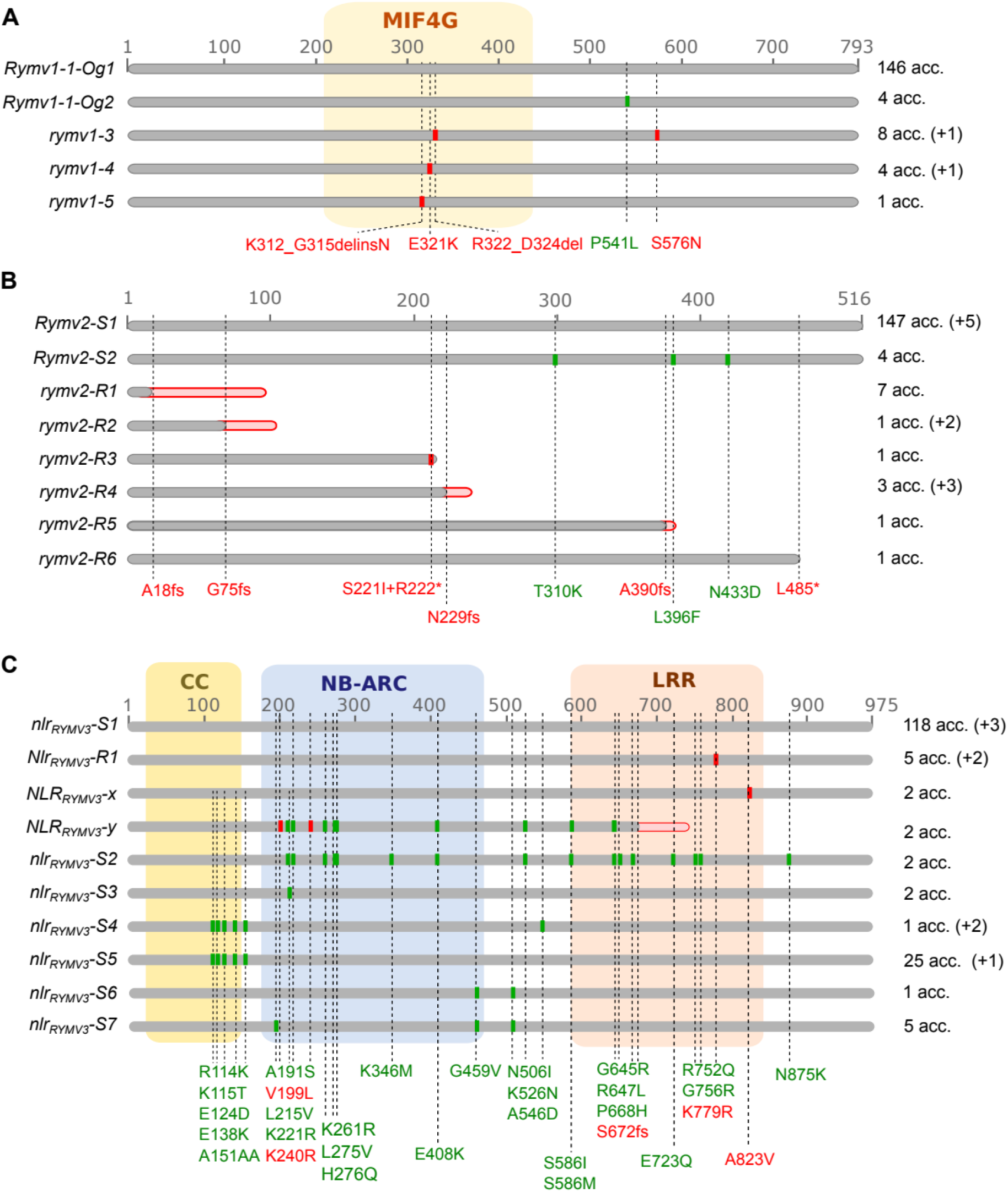
Variants of resistance genes or candidates at the protein level. For *RYMV1* (A), the allele names cited in Albar et al. [14] and Thiemele et al. [12] are used, but an additional protein variant observed in susceptible accessions was given the name “*Rymv1-1-Og2*”, and for greater clarity the allele named “*Rymv1-1-Og*” in [12] was referred to as “*Rymv1-1-Og1*”. For the *RYMV2* (B) and *RYMV3* (C) candidate genes, the different alleles were named according to their association (R) or not (S) with RYMV resistance. The CG14 allele was the reference allele. Polymorphisms associated with resistance are indicated in red, whereas those which are not are in green. Important conserved domains are indicated as colored frames. The number of accessions carrying each allele is indicated on the right of the figure, with a distinction between accessions from the set of 165 fully sequenced accessions (without brackets), and accessions from the nine additional resistant accessions (in brackets). The total number of accessions sometimes differed between genes due to missing data, resulting in undefined alleles.

**Table 2.**
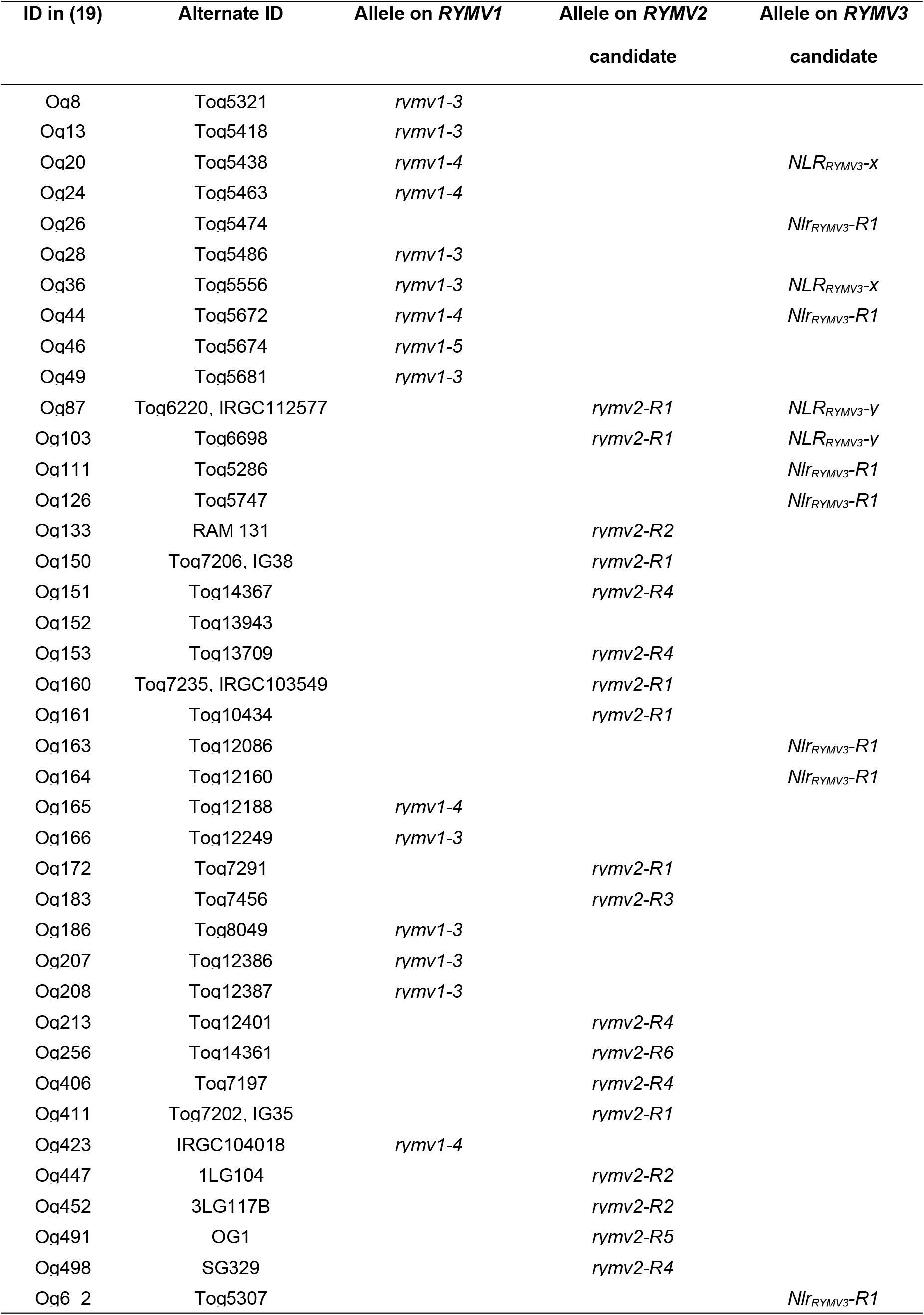
List of the resistant accessions and their alleles on the resistance genes or candidates. Only alleles associated with resistance are indicated.

| ID in (19) | Alternate ID | Allele on *RYMV1* | Allele on *RYMV2*<br>candidate | Allele on *RYMV3*<br>candidate |
| --- | --- | --- | --- | --- |
| Oq8 | Toq5321 | *rymv1-3* |  |  |
| Oq13 | Toq5418 | *rymv1-3* |  |  |
| Oq20 | Toq5438 | *rymv1-4* |  | *NLR<sub>RYMV3</sub>-X* |
| Oq24 | Toq5463 | *rymv1-4* |  |  |
| Oq26 | Toq5474 |  |  | *Nlr<sub>RYMV3</sub>-R1* |
| Oq28 | Toq5486 | *rymv1-3* |  |  |
| Oq36 | Toq5556 | *rymv1-3* |  | *NLR<sub>RYMV3</sub>-X* |
| Oq44 | Toq5672 | *rymv1-4* |  | *Nlr<sub>RYMV3</sub>-R1* |
| Oq46 | Toq5674 | *rymv1-5* |  |  |
| Oq49 | Toq5681 | *rymv1-3* |  |  |
| Oq87 | Toq6220, IRGC112577 |  | *rymv2-R1* | *NLR<sub>RYMV3</sub>-Y* |
| Oq103 | Toq6698 |  | *rymv2-R1* | *NLR<sub>RYMV3</sub>-Y* |
| Oq111 | Toq5286 |  |  | *Nlr<sub>RYMV3</sub>-R1* |
| Oq126 | Toq5747 |  |  | *Nlr<sub>RYMV3</sub>-R1* |
| Oq133 | RAM 131 |  | *rymv2-R2* |  |
| Oq150 | Toq7206, IG38 |  | *rymv2-R1* |  |
| Oq151 | Toq14367 |  | *rymv2-R4* |  |
| Oq152 | Toq13943 |  |  |  |
| Oq153 | Toq13709 |  | *rymv2-R4* |  |
| Oq160 | Toq7235, IRGC103549 |  | *rymv2-R1* |  |
| Oq161 | Toq10434 |  | *rymv2-R1* |  |
| Oq163 | Toq12086 |  |  | *Nlr<sub>RYMV3</sub>-R1* |
| Oq164 | Toq12160 |  |  | *Nlr<sub>RYMV3</sub>-R1* |
| Oq165 | Toq12188 | *rymv1-4* |  |  |
| Oq166 | Toq12249 | *rymv1-3* |  |  |
| Oq172 | Toq7291 |  | *rymv2-R1* |  |
| Oq183 | Toq7456 |  | *rymv2-R3* |  |
| Oq186 | Toq8049 | *rymv1-3* |  |  |
| Oq207 | Toq12386 | *rymv1-3* |  |  |
| Oq208 | Toq12387 | *rymv1-3* |  |  |
| Oq213 | Toq12401 |  | *rymv2-R4* |  |
| Oq256 | Toq14361 |  | *rymv2-R6* |  |
| Oq406 | Toq7197 |  | *rymv2-R4* |  |
| Oq411 | Toq7202, IG35 |  | *rymv2-R1* |  |
| Oq423 | IRGC104018 | *rymv1-4* |  |  |
| Oq447 | 1LG104 |  | *rymv2-R2* |  |
| Oq452 | 3LG117B |  | *rymv2-R2* |  |
| Oq491 | OG1 |  | *rymv2-R5* |  |
| Oq498 | SG329 |  | *rymv2-R4* |  |
| Oq6 2 | Toq5307 |  |  | *Nlr<sub>RYMV3</sub>-R1* |

### Allele mining in *CPR5-1*

In the *CPR5-1* gene, 12 polymorphisms were detected at the nucleotidic level based on genomic data from Cubry et al. [20] (Additional file 1: Table S3). However, the filters used in this analysis hampered detection of the 1 bp-deletion on codon 17 that characterized the allele of the resistant Tog7291 accession [13] because it is located in an artificially created SNP-cluster, probably due to a GCC rich region [23]. Nevertheless, all accessions of the collection had been previously genotyped at this position based on a CAPS marker [13]. Moreover, the deletion was confirmed by manual curation of the read alignment data (BAM file) of the Tog7291 accession. The 13 polymorphisms in the *CPR5-1* gene defined eight haplotypes at the nucleotide level and eight protein variants (Figure 1B; Additional file 1: Table S3). The CG14 reference haplotype was observed in 89% of the accessions. Six haplotypes were characterized by frameshifts (A18fs, G75fs, N229fs, A390fs) or an untimely stop codon (R222*, L485*), leading to truncated forms of the protein, while conserving from 3 to 93% of the protein sequence. Interestingly, these haplotypes concerned 19 accessions that were all highly resistant to RYMV (Table 2; Additional file 1: Table S1), with the most frequent being the Tog7291 haplotype that was previously described in seven accessions [13], while the others were less frequent haplotypes that were found in one to three accessions. Finally, four RYMV-susceptible accessions (Og186, Og426, Og459 and Og89) shared the same haplotype characterized by three SNPs in the introns and three SNPs causing amino-acid substitutions (T310K, L396F, N433D).

**Table 3.** Cosegregation of RYMV resistance and allelic state on *CPR5-1*. F2 plants were evaluated for RYMV resistance based on symptom observations. The phenotype is indicated with R for resistant plants and S for susceptible ones. F2 plants derived from crosses with Tog7291 were not genotyped. For other populations, genotyping on the *CPR5-1* gene was generally performed on all plants with CAPS or dCAPS markers, except for the (Og133 x Og431) population for which genotyping was based on Sanger sequencing and only performed on a subset of 35 plants. The genotype is indicated as “*rymv2-Rx*” for plants homozygous for alleles *rymv2-R2* to *-R6*, “*Rymv1-S1*” for plants homozygous for the *Rymv1-S1* allele, and Htz for heterozygous plants.

| F2 population | Phenotype and genotype |  |  |  |
| --- | --- | --- | --- | --- |
|  | Total | *rymv2-Rx* | Htz | *Rymv2-S1* |
| Og133 (*rymv2-R2*) x Og431 (*Rymv2-S1*) | 11 R, 50S | 10 R | 22 S | 3 S |
| Og133 (*rymv2-R2*) x Tog7291 (*rymv2-R1*) | 55 R, 1S |  |  |  |
| Og183 (*rymv2-R3*) x Og82 (*Rymv2-S1*) | 14 R, 36 S | 14 R | 26 S | 10 S |
| Og213 (*rymv2-R4*) x Og82 (*Rymv2-S1*) | 13 R, 42 S | 13 R | 31 S | 11 S |
| Og213 (*rymv2-R4*) x Tog7291 (*rymv2-R1*) | 100 R |  |  |  |
| Og491 (*rymv2-R5*) x Og431 (*Rymv2-S1*) | 24 R, 47 S | 21 R | 3 R, 30 S | 17 S |
| Og491 (*rymv2-R5*) x Tog7291 (*rymv2-R1*) | 56 R |  |  |  |
| Og256 (*rymv2-R6*) x Og82 (*Rymv2-S1*) | 18 R, 52 S | 18 R, 2S | 37 S | 13 S |
| Og256 (*rymv2-R6*) x Tog7291 (*rymv2-R1*) | 45 R |  |  |  |

### Allele mining in *NLR_RYMV3_*

The variability in *NLR_RYMV3_*, with 66 polymorphisms at the nucleotidic level, was far greater than the variability observed in *RYMV1* and *CPR5-1* (Additional file 1: Table S4). Yet, the polymorphisms identified in the first intron were probably underestimated because of the marked differences between the Nipponbare sequence used as mapping reference and the CG14 sequence, which probably hampered correct mapping and extensive SNP calling in this region. Eleven haplotypes were detected, with the CG14 haplotype being found in 71.5% of the accessions. Forty-nine mutations were located in exons, including 35 that were non-synonymous (Figure 1.C). These mutations defined ten protein variants, three of which were specific to resistant accessions (Table2; Additional file 1: Table S1): two displayed a single amino-acid substitution compared to the reference allele (K779R described in Pidon et al. [18], and A823V), while the third one showed a frameshift in the LRR domain (S672fs) and 11 amino-acid substitutions. The K779R mutation was observed in the two accessions – Tog5307 and Tog5672 – known to carry a resistance allele of *RYMV3* [18], and five that displayed a resistant phenotype but did not carry resistance specific alleles on *RYMV1* or *CPR5-1*, which suggested that their resistance may be associated with the K779R mutation. Conversely, accessions carrying the A823V mutation (Og20, Og36) also had known resistance alleles of *RYMV1* (*rymv1-4* and *rymv1-3*, respectively), and accessions carrying the S672fs mutation (Og87, Og103) had a *CPR5-1* allele associated with resistance.

**Table 4.** Segregation of T-DNA and RYMV resistance in progenies. A pseudo-T3 progeny derived from the 3D-01842 mutant, and F3 progenies derived from the 3A-06612 mutant were analyzed. The phenotype is indicated with “R” for resistance and “S” for susceptibility. The genotype is indicated with “WT” for plants without the T-DNA insertion, “Mut” for plants homozygous for the T-DNA insertion and “Htz” for plants heterozygous at the T-DNA insertion site.

| Mutant | WT | Htz | Mut |
| --- | --- | --- | --- |
| 3D-01842 | 12 S | 34 S | 15 R |
| 3A-06612 | 14 S | 23 S | 18 R, 1S |

Ultimately, all resistant accessions described in the *O. glaberrima* collection carried an allele associated with resistance in at least one of the three analyzed genes (Table 2).

### Comparison with *O. sativa*

Moreover, we looked for polymorphisms at *RYMV1, CPR5-1* and *NLR_RYMV3_* in *O. sativa* based on the SNP-Seek database [24], which pools genotyping data from the 3000 Rice Genomes Project [25]. Seventeen non-synonymous mutations were identified in *RYMV1*. They resulted in amino-acid substitutions or small deletions, but only three occurred in the middle domain of the eukaryotic initiation factor 4G (MIF4G), where all mutations conferring resistance to RYMV were located (Additional file 1: Table S5). One of them (A303D) was present only in the few *O. glaberrima* accessions included in the 3000 Rice Genomes Project, as well as in all accessions from our *O. glaberrima* collection. This mutation was therefore considered to be specific to *O. glaberrima* and not associated with resistance to RYMV. The two others, i.e. K352R and P395S, were detected in four and ten accessions, respectively. While located in the MIF4G domain, they did not occur in the 15 amino-acid region which was mutated in the resistance alleles described so far, but instead were detected at least 28 amino-acids downstream. Twenty-three non-synonymous mutations were detected in the *CPR5-1* gene (Additional file 1: Table S5). However, none of them led to an untimely stop codon or frameshift. Similarly to what we observed in our *O. glaberrima* dataset, *O. sativa* presented high variability at the *NLR_RYMV3_* locus, with 112 non-synonymous mutations (Additional file 1: Table S5). Eight mutations were detected in 10.4% of the accessions and resulted in stop codons or frameshifts. The 104 others were in frame mutations, leading to amino-acid substitutions or single amino acid insertions or deletions in the protein. Interestingly, three *O. sativa* spp. *indica* accessions carried the K779R mutation associated with RYMV resistance in *O. glaberrima*. These accessions shared a very specific haplotype, with 27 additional uncommon non-synonymous mutations that differentiated them from both other *O. sativa* and *O. glaberrima* accessions.

### Loss-of-function mutations in the *CPR5-1* gene confer resistance to RYMV

A genetic analysis was performed to check the association between the truncated CPR5-1 forms identified in *O. glaberrima* and RYMV resistance. Resistant Og256, Og213, Og491, Og133 and Og183 accessions, representing the different truncated forms of CPR5-1, were crossed with a susceptible *O. glaberrima* accession (Og82 or Og431) and with the resistant Tog7291 accession, whose resistance is controlled by *RYMV2* [13]. F2 seeds were obtained for all combinations except (Og183 x Tog7291), and at least 45 F2 plants per population were phenotyped for RYMV resistance. The resistance segregations noted in all populations developed with the susceptible Og82 or Og431 accessions were in agreement with a 1R:3S segregation ratio (Table 3), indicating monogenic and recessive control of resistance. Genotyping on the *CPR5-1* gene was performed on a total of 281 plants based on Sanger sequencing for the Og133-derived population and cleaved amplified polymorphic sequence (CAPS) or derived cleaved amplified polymorphic sequence (dCAPS) markers for all the other populations. Most of the plants homozygous for a truncated form of the protein were resistant (76 out of 79), while most of the others (200 out of 202) were susceptible, showing a close association between the *CPR5-1* allelic state and RYMV resistance. We hypothesized that the five plants that did not fit this pattern were misclassified, presumably because of lack of inoculation or resistance breakdown [17,18,21]. Besides, 256 F2 plants from populations developed with the resistant Tog7291 accession were resistant (Table 3), while a single one was susceptible. These results demonstrated that the different truncated forms of CPR5-1 were resistance alleles of the *RYMV2* recessive resistance gene.

In addition, *O. sativa* lines mutated in the *CPR5-1* gene were characterized. T-DNA insertional mutant lines tagged in the *CPR5-1* gene were identified by searching the flanking sequence database [26] of the mutant library developed by Jeon et al. [27] and Jeong et al. [28]. Two independent T-DNA insertions in the *CPR5-1* gene were confirmed by sequencing the T-DNA flanking regions. In the 3D-01842 line, T-DNA was inserted 1975 bp downstream of the ATG, in the fourth exon; in the 3A-06612 line, T-DNA was inserted 315 bp downstream of the ATG in the first intron (Figure 2A). Phenotyping of these mutants was performed on a minimum of 12 plants homozygous for the insertion. Ten weeks post-sowing, 3D-01842 and 3A-06612 non-inoculated mutants did not show any visible differences in plant morphology or development compared to the wild-type controls (Figure 2B). The mutants inoculated with RYMV did not show any symptoms or growth reduction compared to the non-inoculated controls, while wild-type plants expressed very clear yellowing and mottling symptoms, a marked growth reduction or even growth arrest (Figure 2B and C). In addition, contrary to wild-type plants, mutants did not accumulate the virus according to the ELISA test findings. A total of 117 pseudo-T3 or F3 plants which segregated for one or another T-DNA insertion were analyzed. Except for one plant, a perfect co-segregation was observed between resistance and T-DNA insertions at the homozygous state (Table 4). This indicated that, in both *O. sativa* and *O. glaberrima*, altered forms of CPR5-1 lead to RYMV resistance.

**Figure 2.**
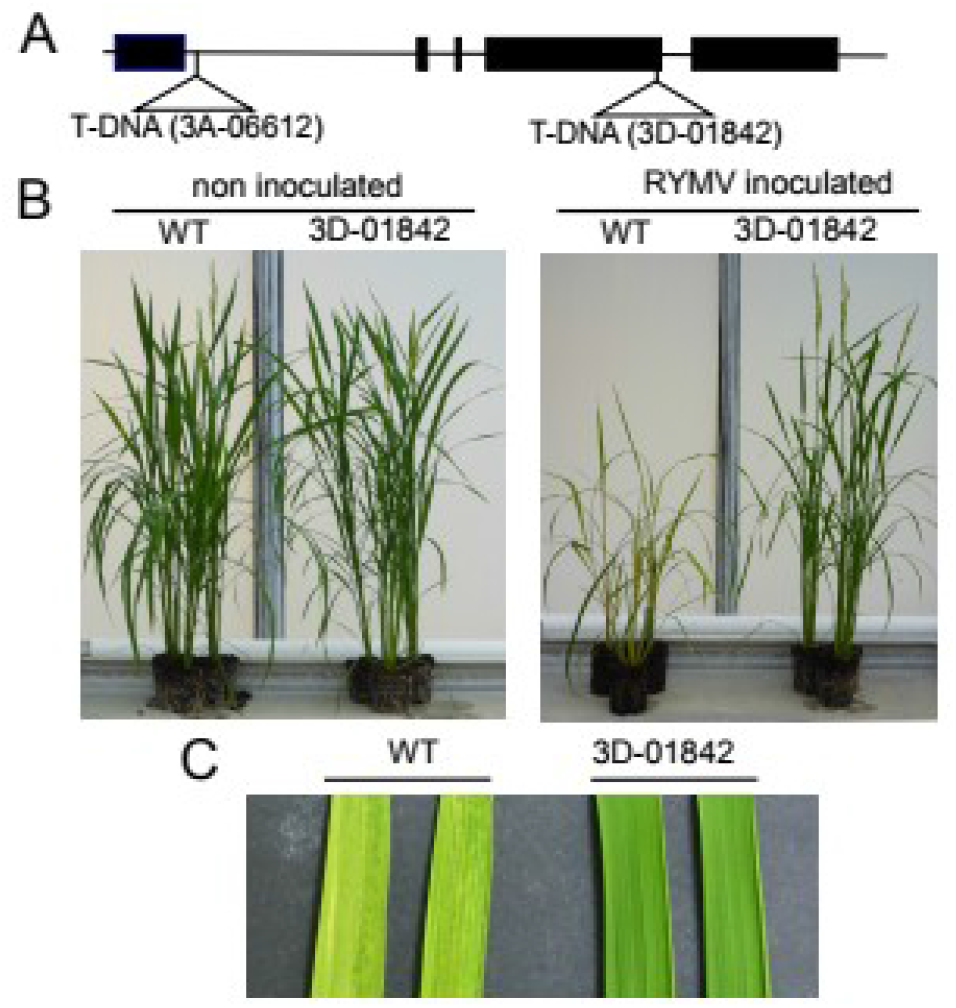
Characterization of T-DNA mutant lines 3D-01842 and 3A-06612. (A) Structure of the *CPR5-1* gene and positions of T-DNA, inserted 315 bp downstream of the ATG in the 3A-06612 line and 1975 bp downstream of the ATG in the 3D-0184 line. Exons are represented as black boxes and introns as black lines. (B) Phenotype of wild-type controls and the 3D-0184 mutant on 10-week old non-inoculated plants, and 8 weeks after RYMV inoculation on inoculated plants, at the whole plant level. (C) Yellowing of leaves of wild-type controls and the 3D-0184 mutant 2 weeks post-inoculation with RYMV.

## Discussion

The results of Orjuela et al. [13] strongly suggested that *CPR5-1* is the *RYMV2* gene, conferring resistance to RYMV. Here we validated this hypothesis using two independent T-DNA mutants in *O. sativa* and six different alleles leading to truncated forms of the protein in *O. glaberrima*. Although *Arabidopsis cpr5* mutants are known to be involved in biotic resistance [15], this is the first time that this gene has been described as a natural resistance gene in a crop species. In *Arabidopsis*, CPR5 is a transmembrane nucleoporin involved in the membrane ring of the nuclear pore complex [16]. Loss of function mutations permeabilize the nuclear pore and mediate the activation of cell cycle transcription factors, leading to defense gene expression. Constitutive resistance to several pathogens is one of the resulting phenotypes, but the mutant shows additional deleterious developmental phenotypes, such as reduced size [15] and seed yield [29], which would be incompatible with breeding strategies for biotic resistance in crops. Some of the six *RYMV2* resistance alleles identified in *O. glaberrima* were very probably null alleles as stop or frameshift mutations were found to occur far upstream, whereas the *rymv2-R6* allele retained 93% of the wild-type protein sequence. Whether the protein completely loses its cellular function or not remains to be investigated. However, based on the homology with *Arabidopsis* [16], even the *rymv2-R6* allele would lack at least one of the transmembrane domains. Unexpectedly, no obvious deleterious phenotype appeared to be associated with these mutations. In addition, the detection of several null alleles that have evolved independently and have been maintained hardly supports a strong deleterious effect of *CPR5-1* knock-out. This could be explained by the presence of two *Arabidopsis CPR5* homologs in rice which may have partial functional redundancy. It is also possible that the functional homolog of *Arabidopsis CPR5* is *CPR5-2* and not *CPR5-1*. The cellular function of each copy will have to be further analyzed. However, our results suggest that the use of null or truncated *CPR5-1* alleles in rice breeding programs, either by introgression from *O. glaberrima* or by mutagenesis, would be an effective way to achieve RYMV resistance. Similar mutations may provide resistance in other pathosystems and allele mining in species that harbor two homologs of *Arabidopsis CPR5*, such as other cereal species, may help uncover new pathways of pathogen resistance.

Contrary to what was observed for accessions carrying *RYMV1* or *RYMV3* resistance alleles, no accessions carrying a *RYMV2* resistance allele showed resistance in 100% of the plants screened. This may have resulted from incomplete resistance or resistance-breaking events. The high rate of resistance-breakdown reported by Pinel-Galzi et al. [17] on the Tog7291 accession carrying the *rymv2-R1* resistance allele suggests that resistance-breaking events is the most likely hypothesis. Indeed, they reported resistance-breaking rates of up to 96% depending on isolates, while other results reported on *RYMV1* [30] and *RYMV3* [18] suggested less frequent resistance-breakdown on those genes.

This study also revealed new resistance sources without a resistance allele at the *RYMV2* locus. The *RYMV1* locus has been the focus of extensive analysis in recent years [12,14,31]. This larger scale study revealed two additional accessions carrying known resistance alleles but did not uncover any new resistance alleles. On the *NLR_RYMV3_* gene, we identified five additional accessions showing the K779R amino-acid substitution in the LRR region that was proposed as being responsible for a high resistance phenotype [18]. These five accessions displayed high resistance to RYMV, which further strengthens the candidate status of the *NLR_RYMV3_* gene, and particularly the K779R mutation, but formal functional validation is still needed to confirm this. Two other haplotypes at *NLR_RYMV3_* were specific to resistant accessions but the corresponding accessions carried alleles on *RYMV1* and *RYMV2*, which would be sufficient to explain their high resistance level. Furthermore, the *NLR_RYMV3_-y* sequence variant was characterized by a truncated LRR domain, suggesting a loss of function, which is not consistent with a gain of resistance. We think it is likely that those two sequence variants do not confer resistance to RYMV but further genetic analyses would be necessary to confirm this.

In contrast, we did not find any convincing candidate resistance alleles on *RYMV1* and *RYMV2* genes among accessions from the 3000 Rice Genomes Project [25], which mostly includes *O. sativa* accessions. At the *RYMV1* locus, two rare mutations were identified in the MIF4G domain and would require further analysis. However, based on the predicted 3D structure of the MIF4G domain [14], they occurred downstream of the α-helical hairpin that forms a protrusion where mutations known to be responsible for high resistance are located. We therefore do not consider these mutations as likely candidates for resistance. High variability was observed at the *NLR_RYMV3_* locus and a simple sequence analysis would not be sufficient to pinpoint mutations that may be involved in resistance. In particular, the K779R mutation, which is associated with resistance in *O. glaberrima,* has been detected in three *O. sativa* spp. *indica* accessions. However it is hard to speculate on their resistance, as these accessions were also characterized by 14 additional rare mutations in the LRR domain.

The probable absence of candidate resistance alleles on *RYMV1* and *RYMV2* within accessions of the 3000 Rice Genomes Project—mainly *O. sativa*, as mentioned above—is in agreement with the scant resistance observed in this species based on phenotypic screening. Indeed, only two accessions with a high level of resistance to RYMV, like those described in this study, have been reported in *O. sativa* [10, 11]. These two accessions, originating from East Africa, both carry the *rymv1-2* resistance allele. This result contrasted with the relatively high number of resistance alleles detected in *O. glaberrima*. Out of the 268 accessions of the collection used in this study, 40 highly resistant accessions were detected, which corresponded to approximately 15% of the collection. Yet this rate was probably overestimated because about ten accessions previously identified as resistant [12, 32] were deliberately included when the collection was set up [19]. The actual rate of resistant accessions in *O. glaberrima* is probably closer to 8%, which is the rate calculated on the basis of the 148 accessions newly evaluated in this study and for which we did not have any *a priori* knowledge. Still, this rate is very much higher than in *O. sativa*.

The diversity profiles on *RYMV1*, *RYMV2* and *NLR_RYMV3_* genes were contrasted. First, we observed a high number of mutations at the *NLR_RYMV3_* gene, with 35 non-synonymous mutations detected in the *O. glaberrima* collection. Such high variability was expected and has been widely documented for the *NLR* gene family, which is known to be hypermutagenic and frequently under balancing or diversifying selection as a result of the arms race between plants and pathogens [33–38]. Secondly, *RYMV1* and *RYMV2* presented lower variability, with five and ten non-synonymous mutations detected, respectively, in accordance with their central role in plant cells. Indeed, *RYMV1* codes for eIF(iso)4G-1 [14], a translation initiation factor that is part of the cell translation machinery, while the *Arabidopsis* gene homologous to *RYMV2* codes for a component of the nuclear pore complex. These two genes are therefore assumed to be under conservative selection. Interestingly, three out of five non-synonymous mutations in *RYMV1* and six out of ten in *RYMV2* were directly involved in the resistance phenotype. In a similar gene/pathogen interaction, the results of Charron et al. [39] provided evidence of diversifying selection on the eIF4E locus that would at least partially be driven by potyvirus-induced selection pressure. As RYMV emerged quite recently, in the mid-19th century [40, 41], there has not been a long co-evolution between the virus and *O. glaberrima* that could have explained the allelic diversity observed at the resistance loci. However, selection pressure on these loci may have been exerted by other viruses using these exact plant factors.

The different resistance genes and alleles were positioned on the genetic diversity tree of the species proposed by Orjuela et al. [19] and on a map according to the geographical origin of the accessions. For all three resistance genes, accessions carrying the same resistance allele were generally showing a similar geographic origin (Figure 3) and were clustered on the genetic diversity tree (Additional file 2: Figure S1), as expected since *O. glaberrima* has geographically-based population structuring [42]. More surprisingly, accessions with different *RYMV1* or *RYMV2* resistance alleles also appeared to be clustered, despite the independence of the mutations characterizing the alleles. Accessions with *RYMV2* resistance alleles were located west of the Benin-Niger axis, while accessions with *RYMV1* resistance alleles were located east of this axis. *RYMV3* apparently did not fit this distribution pattern, but the low number of accessions limited the scope of these findings. Several hypotheses may explain the observed *RYMV1* and *RYMV2* structuring. First, both eIF(iso)4G and CPR5-1 – if confirmed as a nucleoporin – are part of large protein complexes. Their variability may have been driven by the genetic structuring of other members of the same complexes. Besides, the environmental conditions may have led to a difference in selection pressure on *RYMV1* and *RYMV2,* or members of the complex in which they participate. Accessions with *RYMV2* resistance alleles mainly originated from regions of historically dense rice cultivation, while accessions with *RYMV1* resistance alleles originated from regions where rice was more sparsely cultivated, according to Portères [43]. This axis is also compatible with the separation of two distinct genetic groups of RYMV isolates [40, 41], one of which – the easternmost – includes hypervirulent isolates able to overcome most known resistance sources [30]. As underlined previously, the diversification of resistance genes under a selection pressure exerted by RYMV is quite improbable, but the opposite hypothesis has yet to be investigated, along with the impact of environmental conditions on both the virus and resistance gene diversity. Moreover, like *A. thaliana CPR5* [16], if *RYMV2* is a regulator of effector-triggered immunity and programmed cell death, it may confer resistance to several pathogens and could have evolved under selection pressure exerted by another pathogen. We would be unable to perform a more detailed population genetics analysis due to the limited number of resistant accessions available, but additional collections have been described [42, 44] and should now be investigated.

**Figure 3.**
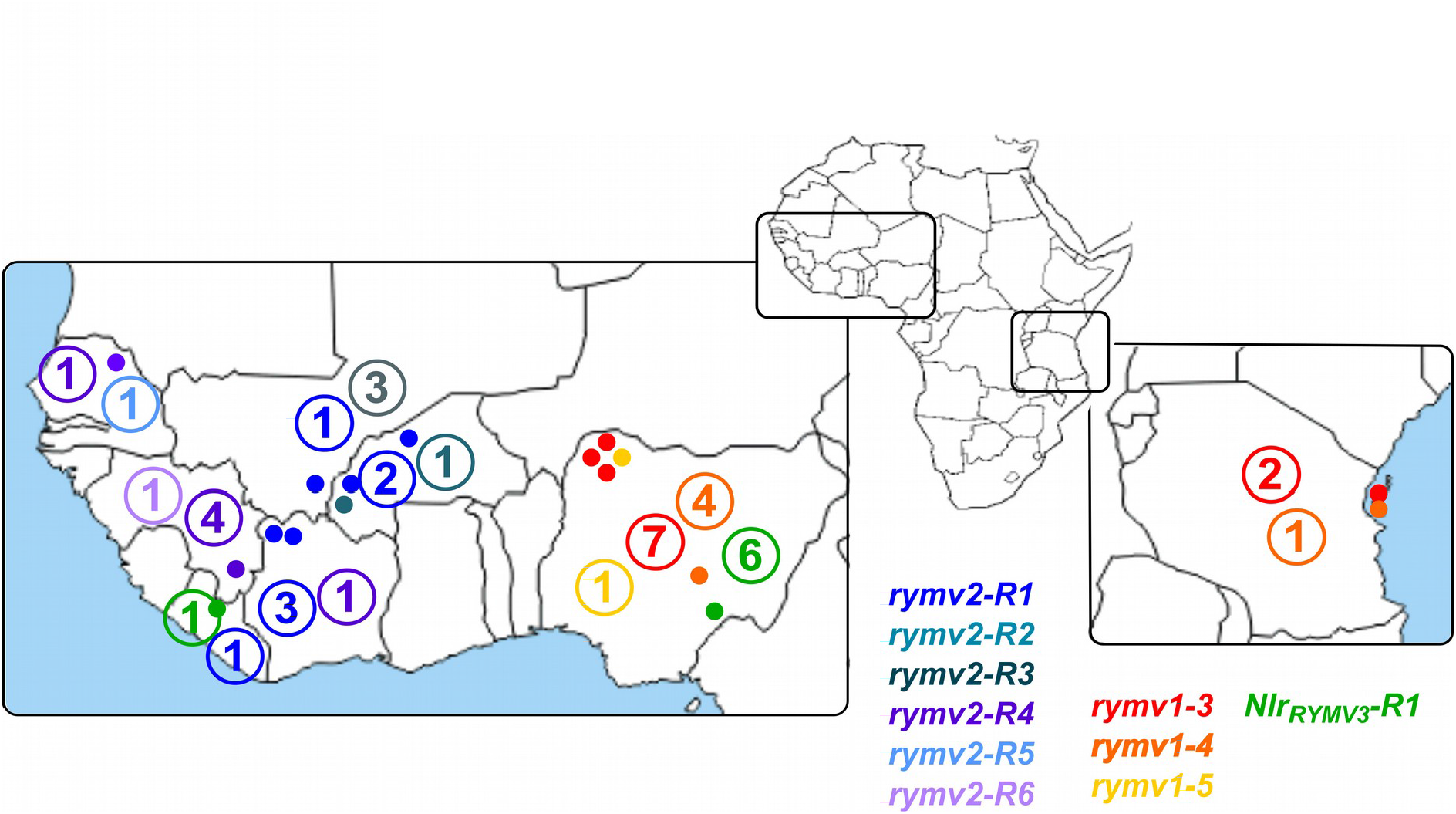
Geographical origin of accessions carrying resistance alleles on *RYMV1*, *RYMV2* and *NLR_RYMV3_* genes. The geographical origins of the accessions were obtained from Cubry et al. [20]. Accessions for which GPS coordinates were available are represented by colored points. In each country, the total number of accessions carrying a specific allele (with or without GPS coordinates) is indicated as a number.

## Conclusions

Our results highlighted the allelic diversity in the three known resistance genes against RYMV. All 40 *O. glaberrima* accessions identified as being highly resistant in this study carried at least one of the confirmed or candidate resistance alleles on *RYMV1*, *RYMV2* and *NLR_RYMV3_* (Table 2). This suggests that we have probably identified most of the resistance genes that occur in *O. glaberrima*, even though wild relative species, such as *Oryza barthii,* may also contain original resistance sources. Sound knowledge on resistance genes against RYMV and their diversity is thus now available, as well as a good assessment of the frequency and molecular determinants of resistance-breakdown in controlled conditions [17,21,30]. This knowledge provides an opportunity to design strategies of resistance gene deployment that will optimize resistance durability. Previous results suggest that all three genes are effective against a large spectrum of RYMV isolates. However, the high capacity of some virus isolates to evolve and overcome resistance prompts breeders to manage the use of resistance genes by pyramiding, variety mixtures or temporal rotation, depending on the local virus diversity. So far, a single *RYMV1* resistance allele originating from *O. sativa* accessions has been transferred into high-yielding varieties [45, 46] that are about to be deployed in the field. The use of additional resistance alleles or genes, and combinations, should be promoted to increase RYMV resistance sustainability.

## Methods

### Plant material

The *O. glaberrima* collection used in this study was described in Orjuela et al. [19]. This collection was jointly established by the French National Research Institute for Sustainable Development (IRD) and the Africa Rice Center and the accessions studied were selected for their current breeding impact and geographical distribution. Accessions identifiers (ID) of Orjuela et al. [19] are generally used, except for the CG14 reference accession or accessions previously characterized for RYMV resistance for which the previously used names were adopted [12, 13]. Correspondences between the different ID are given in Additional file 1: Table S1.

T-DNA mutant lines were obtained from the Pohang University of Science and Technology, Pohang, Korea [27, 28]. The 3D-01842 line was derived from the Hwayoung variety and the 3A-06612 line from the Dongjin variety. For the 3D-01842 line, a plant heterozygous for the T-DNA insertion in the *CPR5-1* gene was self-pollinated to develop pseudo-T2 and pseudo-T3 progenies that segregated for the insertion. As no plant heterozygous for the T-DNA insertion was available for the 3A-06612 line, a plant homozygous for the insertion was crossed with the Dongjin variety and F1 hybrids were selfed to successively derive F2, and F3 progenies that segregated for the T-DNA insertion.

### Resistance evaluation

Plants were grown in greenhouses and mechanically inoculated about 2 weeks after sowing with an RYMV isolate originating from Burkina Faso (BF1). The resistance evaluations were based on symptom observation and confirmed when necessary with DAS-ELISA to estimate the virus content in leaf samples harvested 2-3 weeks post-inoculation. Details on t he inoculation and ELISA protocols were previously described in Pinel-Galzi et al. [47].

### Genomic data

Genomic data on *O. glaberrima* accessions based on high coverage genomic sequencing (average 35X, range 20-55X) were obtained from Cubry et al. [20]. The IRGSP 1.0 Nipponbare sequence [48] had been used as reference for mapping and SNP calling. The polymorphism database [49, 50] included genomic data of 163 *O. glaberrima* and 83 *O. barthii* accessions. In a first step, the database was filtered for the 163 *O. glaberrima* accessions and for the ATG-stop codon regions of target genes. The ORGLA04G0147000.1, ORGLA01G0359000.1 and ORGLA11G0175800.1 gene models established on the CG14 accession [1], corresponding to Os04g42140.1, Os01g68970.1 and Os11g43700.1 gene models on the reference Nipponbare sequence, were considered for *RYMV1*, *CPR5-1* and *NLR_RYMV3_*, respectively. The target regions corresponded to positions 24,946,655-24,952,068 on chromosome 4 of the reference Nipponbare sequence for *RYMV1*, 40,071,092-40,073,727 on Nipponbare chromosome 1 for *CPR5-1* and 26,377,263-26,380,577 on Nipponbare chromosome 11 for *NLR_RYMV3_*. Only SNPs and indels that were polymorphic within *O. glaberrima* accessions were conserved. In a second step, polymorphisms were filtered with GATK 3.7 VariantFiltration [51] using the following filters: QUAL<200, MQ0 >4 && MQ0/DP>0.1, DP<10, clusterSize 3 in clusterWindowSize 10, DP>20000. SNPs with more than 10% missing data or heterozygous in more than 10% accessions were filtered out, and genotypes defined based on a single read were considered as missing data. Genomic data of the *O. glaberrima* CG14 [1] and Tog5681 accessions [22] were included in the dataset. The haplotype of the RYMV-susceptible CG14 accession was used as a reference to describe variants observed in the population.

Genomic data from the 3000 Rice Genomes Project [25] were retrieved from the Rice SNP-Seek database [24]. The database was filtered on the target regions indicated above for indels and non-synonymous SNPs from the base SNP set, which includes about 18 million SNPs. SNP effects were retrieved from the database, while indel effects were manually estimated.

The sequence variant nomenclature proposed by Den Dunnen et al. [52] was used to describe the mutations and their effects on CDS and proteins. Based on the results described in this paper or previously [12,14,18], dominant alleles were indicated with the first letter in upper case and recessive ones with the first letter in lower case; when there was no preferred hypothesis, allele names were written with all letters in upper case. For the *RYMV2* and *RYMV3* candidate genes, the different alleles were named depending on their association (R) or not (S) with RYMV resistance.

### Sanger sequencing and molecular markers

PCR amplifications were performed on leaf extracts or DNA, as described in Orjuela et al. [13]. Primers were designed using Primer3 [53], except for the primers used for dCAPS markers, which were designed with the dCAPS Finder tool [54]. Primer sequences are provided in the Additional file 2: Figure S2. Partial or complete Sanger sequencing of *RYMV1*, *CPR5-1* and *NLR_RYMV3_* genes was performed on PCR amplification fragments and subcontracted to Genewiz (Takeley, UK).

CAPS and dCAPS markers were designed on the *CPR5-1* gene to genotype polymorphic loci identified in *O. glaberrima* accessions. Marker characteristics are described in the Additional file 2: Figure S2 and Table S6.

The T-DNA segregation analysis was based on the amplification of a T-DNA-specific fragment and a gene-specific fragment involving a common primer. Primer sequences and positions are indicated in the Additional file 2: Figure S2.

## List of abbreviations

CPR5: constitutive expression of pathogenesis-related protein-5
DAS-ELISA: double antibody sandwich-enzyme-linked immunosorbent assay
dCAPS: derived cleaved amplified polymorphic sequence
ID: identifier
Indel: small insertion/deletion
MIF4G: middle domain of eukaryotic initiation factor 4G
LRR: leucine-rich repeat
NLR: nucleotide-binding domain and leucine-rich repeat containing domain
PCR: polymerase chain reaction
RYMV: *Rice yellow mosaic virus*
SNP: single nucleotide polymorphism

## Declarations

### Ethics approval and consent to participate

Not applicable

### Consent for publication

Not applicable

### Availability of data and materials

The *O. glaberrima* genomic dataset analysed in the current study are available in the IRD Gigwa repository, https://gigwa.ird.fr/gigwa/. All additional data generated in this study are included in the present article and its supplementary information files.

### Authors’ contributions

AG, HP and LA planned and designed the experiments, SC developed the populations, SC, HP and LA performed phenotyping, genotyping and sequencing, HP and LA analyzed the data and wrote the paper, AG and SC revised the paper.

## Acknowledgements

We thank Christine Dubreuil-Tranchant and François Sabot for their help in the bioinformatics analysis, Gatean Maillot for technical help in T-DNA mutant characterization and Harold Chrestin for seed multiplication. We are also grateful to Denis Fargette, François Anthony, Mathias Lorieux and Nils Poulicard for helpful discussions. We thank Gynheung An for providing the T-DNA mutant lines.

## Additional file captions

**Additional file 1 (.xls)**

**Additional file 1: Table S1. ID, phenotype and genotype of accessions characterized for RYMV resistance.** Resistance to RYMV was evaluated after mechanical inoculation of the BF1 isolate in this study or in previous studies [12, 13]. Alleles on resistance genes or candidates refer to the results presented in the Additional File 1: Table S2, Table S3, Table S4 or in previous studies [12].

**Additional file 1: Table S2. Genotype on the *RYMV1* resistance gene.** Only positions where polymorphisms were detected in the *O. glaberrima* collection analyzed in Cubry et al. [20] were included. Nucleotide positions refer to the IRGSP1.0 reference sequence of the *O. sativa* Nipponbare accession [46] that was used as mapping reference. The effect of the mutations are based on the ORGLA04G0147000.1 gene model established on the *O. glaberrima* CG14 accession [1]. Mutations are described according to the nomenclature proposed by Den Dunnen et al. [48], except that synonymous mutations and mutations occurring in an intron are denoted “syn” and “intron”, respectively. Different variants at the protein level were considered as different alleles. Names for resistance alleles were previously attributed by Albar et al. [14] and Thiemele et al. [12], but an additional protein variant observed in susceptible accessions was given the name “*Rymv1-1-Og2*”, and for greater clarity the allele named “Rymv1-1-Og” in [12] was referred to as “*Rymv1-1-Og1*”.

**Additional file 1: Table S3. Genotype on the *CPR5-1* gene, candidate for *RYMV2*.** Only positions were polymorphisms were detected in the *O. glaberrima* collection analyzed in Cubry et al. [20] were included. Nucleotide positions referred to the IRGSP1.0 reference sequence of the *O. sativa* Nipponbare accession [46] that was used as mapping reference. The effects of the mutations are based on the ORGLA01G0359000.1 gene model established on the *O. glaberrima* CG14 accession [1]. Mutations are described according to the nomenclature proposed by Den Dunnen et al. [48], except that synonymous mutations and mutations occurring in an intron are noted “syn” and “intron”, respectively. Different variants at the protein level were considered as different alleles. The allele names were chosen to distinguish protein variants associated or not with RYMV resistance.

**Additional file 1: Table S4. Genotype on the *NLR_RYMV3_* gene, candidate for *RYMV3*.** Only positions were polymorphisms were detected in to the *O. glaberrima* collection analyzed in Cubry et al. (2018) were included. Nucleotide positions refer to the IRGSP1.0 reference sequence of the *O. sativa* Nipponbare accession [46] that was used as mapping reference. The effects of the mutations are based on the ORGLA11G0175800.1 gene model established on the *O. glaberrima* CG14 accession (Wang et al., 2014). Mutations are described according to the nomenclature proposed by Den Dunnen et al. [48], except that synonymous mutations and mutations occurring in an intron are noted “syn” and “intron”, respectively. Different variants at the protein level were considered as different alleles. The allele names were chosen to distinguish protein variants associated or not with RYMV resistance.

**Additional file 1: Table S5. Diversity on RYMV resistance genes or candidates in accessions from the 3000 Rice Genomes Project** [25]. Only non-synonymous SNPs from the base SNPs set are reported here. SNP effects were retrieved from the SNP-Seek database [24] and indels effects were evaluated manually. The effects of mutations on CDS and proteins are based on the Os04g42140.1 and Os01g68970.1 gene models established on the Nipponbare IRGSP1.0 sequence [46], for *RYMV1* and *CPR5-1,* respectively. For *NLR_RYMV3_,* the CDS is based on the Os11g43700.1 gene mode, except that the ATG codon was shifted from 180 nucleotides downstream of the original starting codon to best fit the corresponding CDS of the ORGLA11G0175800.1 gene model established on CG14 reference sequence. Effects on the CDS and protein were thus adapted. Frequency refers to the percentage of the alternate variant in the complete set of accessions. Mutations located in the PFAM domains MA3, MIF4G and LRR and in the HMM Panther hit LRR are indicated.

**Additional file 2 (.pdf)**

**Additional file 2: Figure S1.** Positions of accessions with resistance alleles of RYMV1, RYMV2 and NLRRYMV3 genes on the genetic diversity tree. Susceptible accessions are colored in dark grey and accessions not evaluated for resistance in light grey. Adapted from the genetic tree of Orjuela et al. [19].

**Additional file 2: Figure S2.** Characteristics of primers and amplified fragments for markers or Sanger sequencing. Genes are represented as grey boxes for exons and grey lines for introns. Primers are represented as triangles and the numbers below the triangle refer to the corresponding sequences. (a, b, c) Blue traits represent fragments that were amplified and then sequenced with the primers colored in red. (c) Amplification fragments corresponding to the CAPS or dCAPS markers designed on the CPR5-1 gene are represented as green traits. Additional information on these markers is provided in Additional file 2: Table S6. (d) Position of T-DNA insertions in the CPR5-1 gene in lines 3A-06612 and 3D-01842 are indicated. The T-DNA-specific and gene-specific primers used for sequencing the T-DNA flanking site and genotyping for the presence/absence of insertions are indicated in brown and blue, respectively.

**Additional file 2: Table S6.** Characteristics of CAPS and dCAPS markers. Marker names indicate whether there are CAPS or dCAPS markers and which alleles of the CPR5-1 gene they target. The bracketed number before the primer sequences refer to the reference of primers in Additional file 2: Figure S2. The size of the fragments expected in plants with the reference haplotype of CG14 (WT) or the alternate haplotypes (R) are indicated, except fragments below 30 bp that are uneasily detected by agarose electrophoresis. The CAPS-CPR5-1-R1 marker had already been described in Orjuela et al. [13].

