## Additional file 2 for "Allele mining unlocks the identification of RYMV resistance genes and alleles in African cultivated rice"

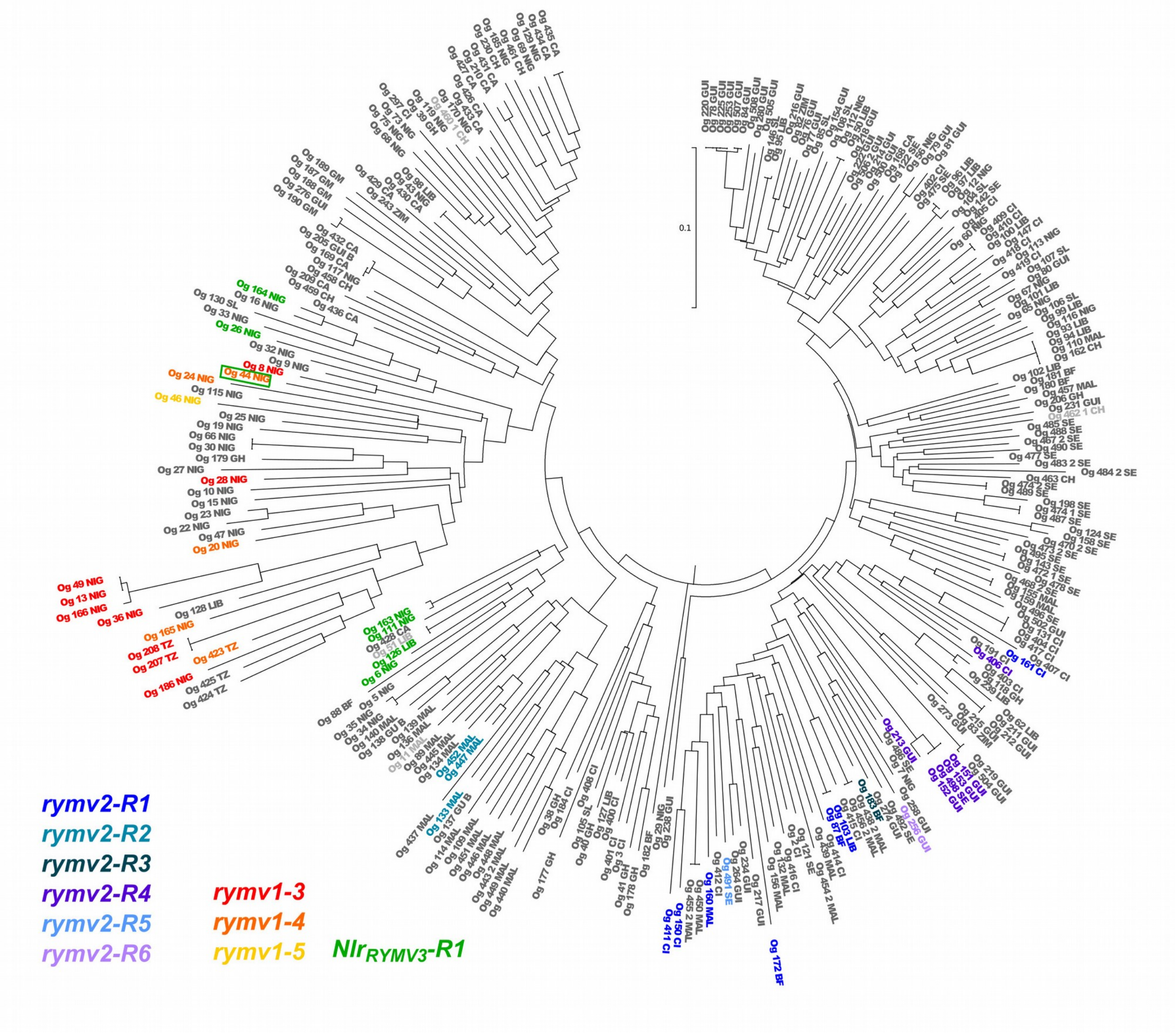


**Figure S1. Positions of accessions with resistance alleles of *RYMV1*, *RYMV2* and *NLR$_{RYMV3}$* genes on the genetic diversity tree.** Susceptible accessions are colored in dark grey and accessions not evaluated for resistance in light grey. Adapted from the genetic tree of Orjuela et al. [19].

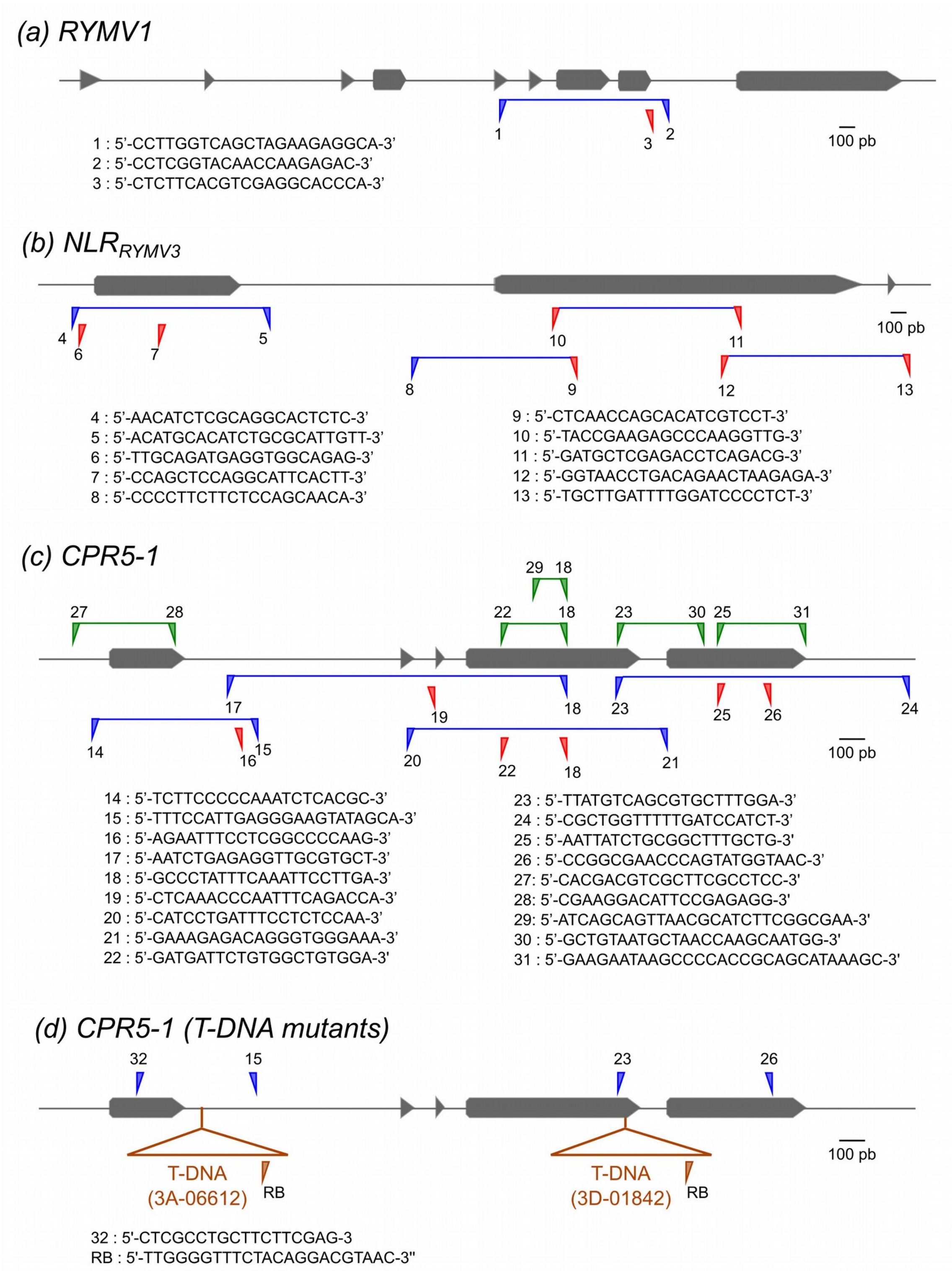


**Figure S2. Characteristics of primers and amplified fragments for markers or Sanger sequencing.** Genes are represented as grey boxes for exons and grey lines for introns. Primers are represented as triangles and the numbers below the triangle refer to the corresponding sequences. (a, b, c) Blue traits represent fragments that were amplified and then sequenced with the primers colored in red. (c) Amplification fragments corresponding to the CAPS or dCAPS markers designed on the *CPR5-1* gene are represented as green traits. Additional information on these markers is provided in Additional file 2: Table S6. (d) Position of T-DNA insertions in the *CPR5-1* gene in lines 3A-06612 and 3D-01842 are indicated. The T-DNA-specific and gene-specific primers used for sequencing the T-DNA flanking site and genotyping for the presence/absence of insertions are indicated in brown and blue, respectively.

**Table S6. Characteristics of CAPS and dCAPS markers.** Marker names indicate whether there are CAPS or dCAPS markers and which alleles of the *CPR5-1* gene they target. The bracketed number before the primer sequences refer to the reference of primers in Additional file 2: Figure S2. The size of the fragments expected in plants with the reference haplotype of CG14 (WT) or the alternate haplotypes (R) are indicated, except fragments below 30 bp that are uneasily detected by agarose electrophoresis. The CAPS-CPR5-1-R1 marker had already been described in Orjuela et al. [13].

| Marker | Primers | Restriction Enzyme | Fragment size (bp) |
|---|---|---|---|
| CAPS-CPR5-1-R1 | [27] 5'-CACGACGTCGCTTCGCCTCC-3'<br>[28] 5'-CGAAGGACATTCCGAGAGG-3' | Cfo I | R: 151 + 121 + 71<br>WT: 223 + 121 |
| dCAPS-CPR5-1-R3 | [29] 5'-ATCAGCAGTTAACGCATCTTCGGCGAA-3'<br>[18] 5'-GATGATTCTGTGGCTGTGGA-3' | Hinf I | R: 102<br>WT: 129 |
| CAPS-CPR5-1-R4 | [22] 5'-GATGATTCTGTGGCTGTGGA-3'<br>[18] 5'-GCCCTATTTCAAATTCCTTGA-3' | Eco RV | R: 168 + 78<br>WT: 247 |
| dCAPS-CPR5-1-R5 | [23] 5'-TTATGTCAGCGTGCTTTGGA-3'<br>[30] 5'-GCTGTAATGCTAACCAAGCAATGG-3' | Hae III | R: 329<br>WT: 305 |
| dCAPS-CPR5-1-R6 | [25] 5'-AATTATCTGCGGCTTTGCTG-3'<br>[31] 5'-GAAGAATAAGCCCCACCGCAGCATAAAGC-3' | Alu I | R: 208<br>WT: 236 |
